# MeFSAT: A curated natural product database specific to secondary metabolites of medicinal fungi

**DOI:** 10.1101/2020.12.04.412502

**Authors:** R.P. Vivek-Ananth, Ajaya Kumar Sahoo, Kavyaa Kumaravel, Karthikeyan Mohanraj, Areejit Samal

**Author notes:** Corresponding author (A. Samal). R.P.V., A.K.S. and K.K. contributed equally to this work and should be considered as Joint-First authors.

## Abstract

Fungi are a rich source of secondary metabolites which constitutes a valuable and diverse chemical space of natural products. Medicinal fungi have been used in traditional medicine to treat human ailments for centuries. To date, there is no devoted resource on secondary metabolites and therapeutic uses of medicinal fungi. Such a dedicated resource compiling dispersed information on medicinal fungi across published literature will facilitate ongoing efforts towards natural product based drug discovery. Here, we present the first comprehensive manually curated database on **Me**dicinal **F**ungi **S**econdary metabolites **A**nd **T**herapeutics (MeFSAT) that compiles information on 184 medicinal fungi, 1830 secondary metabolites and 149 therapeutics uses. Importantly, MeFSAT contains a non-redundant *in silico* natural product library of 1830 secondary metabolites along with information on their chemical structures, computed physicochemical properties, drug-likeness properties, predicted ADMET properties, molecular descriptors and predicted human target proteins. By comparing the physicochemical properties of secondary metabolites in MeFSAT with other small molecules collections, we find that fungal secondary metabolites have high stereochemical complexity and shape complexity similar to other natural product libraries. Based on multiple scoring schemes, we have filtered a subset of 228 drug-like secondary metabolites in MeFSAT database. By constructing and analyzing chemical similarity networks, we show that the chemical space of secondary metabolites in MeFSAT is highly diverse. The compiled information in MeFSAT database is openly accessible at: https://cb.imsc.res.in/mefsat/.

## Introduction

Fungi are present in every ecological niche, and thus, face challenge from myriad biotic and abiotic stressors^1,2^. Investigation of the fungal habitats has shed important insights on the stimulation and the biosynthesis of valuable secondary metabolites produced by fungi^3^. As a rich source of secondary metabolites, fungi are valuable contributors to the chemical diversity of the natural product space. Importantly, the fungal secondary metabolome is enriched in bioactive molecules and has immense potential for drug discovery^4^ including antibiotics. Notably, the first broad-spectrum antibiotic, Penicillin, discovered by Alexander Fleming is a fungal secondary metabolite.

Secondary metabolites are small organic compounds mainly produced by plants, fungi and bacteria, and these metabolites are not essential for the growth and reproduction of the organism^5^. As a rich source of bioactive molecules, it is not surprising that natural products have played an indomitable role in the history of drug discovery^6^. By one estimate^6^, natural products have contributed to the discovery of ∼35% drugs approved by the US Food and Drug Administration (FDA) to date. Therefore, significant research effort has been devoted towards development of natural product databases^7–15^ to facilitate computational approaches to drug discovery.

In this context, several natural product databases^8,9,11–14,16^ dedicated to secondary metabolites or phytochemicals of medicinal plants have been built to date. For instance, some of us have built the IMPPAT^14^ database which is the largest dedicated resource on phytochemicals of Indian medicinal plants to date. During manual curation of the IMPPAT^14^ database, we realized that there is no dedicated resource on secondary metabolites of medicinal fungi (or mushrooms) to date. This is surprising given medicinal mushrooms^17–21^ have also been used for centuries in traditional medicine, especially traditional Chinese medicine, to treat human ailments. Therefore, a comprehensive and dedicated resource on secondary metabolites of medicinal fungi along with their therapeutic uses will serve ongoing research efforts towards natural product based drug discovery. Here, we have addressed this unmet need by building a natural product database dedicated to secondary metabolites of medicinal fungi.

The fungal kingdom is very large and encompasses diverse organisms ranging from simple yeasts to mushrooms. Some fungi are considered ‘medicinal’ due to the beneficial bioactivity of their secondary metabolites and/or their usage in systems of traditional medicine to treat human ailments^17–21^. Mushrooms are macrofungi with fruiting bodies^22^ that have been used as food and/or medicine for centuries across many civilizations^17–21,23^. Mushroom-derived preparations have been used in traditional medicine practiced in many Asian countries^24^. Presently, the valuable information on secondary metabolites and therapeutic uses of medicinal fungi is dispersed across published literature including articles and books^17–21^, and this limits its effective use for drug discovery. Moreover, existing microbial natural product databases such as NPATLAS^15^ have certain limitations; they are neither specific to fungi nor do they capture both secondary metabolite and therapeutic use information for medicinal fungi. In other words, a common repository of high-quality information on secondary metabolites and therapeutic uses of medicinal fungi is needed to harness the potential of this chemical space for drug discovery.

In this direction, we present a manually curated database, **Me**dicinal **F**ungi **S**econdary metabolites **A**nd **T**herapeutics (MeFSAT), dedicated to secondary metabolites and therapeutic uses of medicinal fungi (Fig. 1). Briefly, MeFSAT compiles manually curated information on 184 medicinal fungi, 1830 secondary metabolites and 149 therapeutic uses from published literature. Importantly, we have created a non-redundant library of 1830 fungal secondary metabolites with information on their chemical structure. For the 1830 fungal secondary metabolites, we have compiled information on their physicochemical properties, drug-likeness properties, predicted ADMET properties and molecular descriptors using cheminformatics tools^25–29^. Further, we have gathered information on the predicted human target proteins of the secondary metabolites from the STITCH database^30^. After constructing the MeFSAT database, we have filtered a subset of 228 drug-like fungal secondary metabolites based on six drug-likeness scoring schemes. Moreover, we have also compared the physicochemical properties of the secondary metabolites in MeFSAT database with those of small molecules in three other libraries. Lastly, we have constructed and analyzed two chemical similarity networks (CSNs) corresponding to the 1830 secondary metabolites and the 228 drug-like secondary metabolites in MeFSAT database, and these networks highlight the diverse chemical space of secondary metabolites produced by medicinal fungi. The compiled and curated information in the MeFSAT database is openly accessible at: https://cb.imsc.res.in/mefsat/.

**Figure 1.**
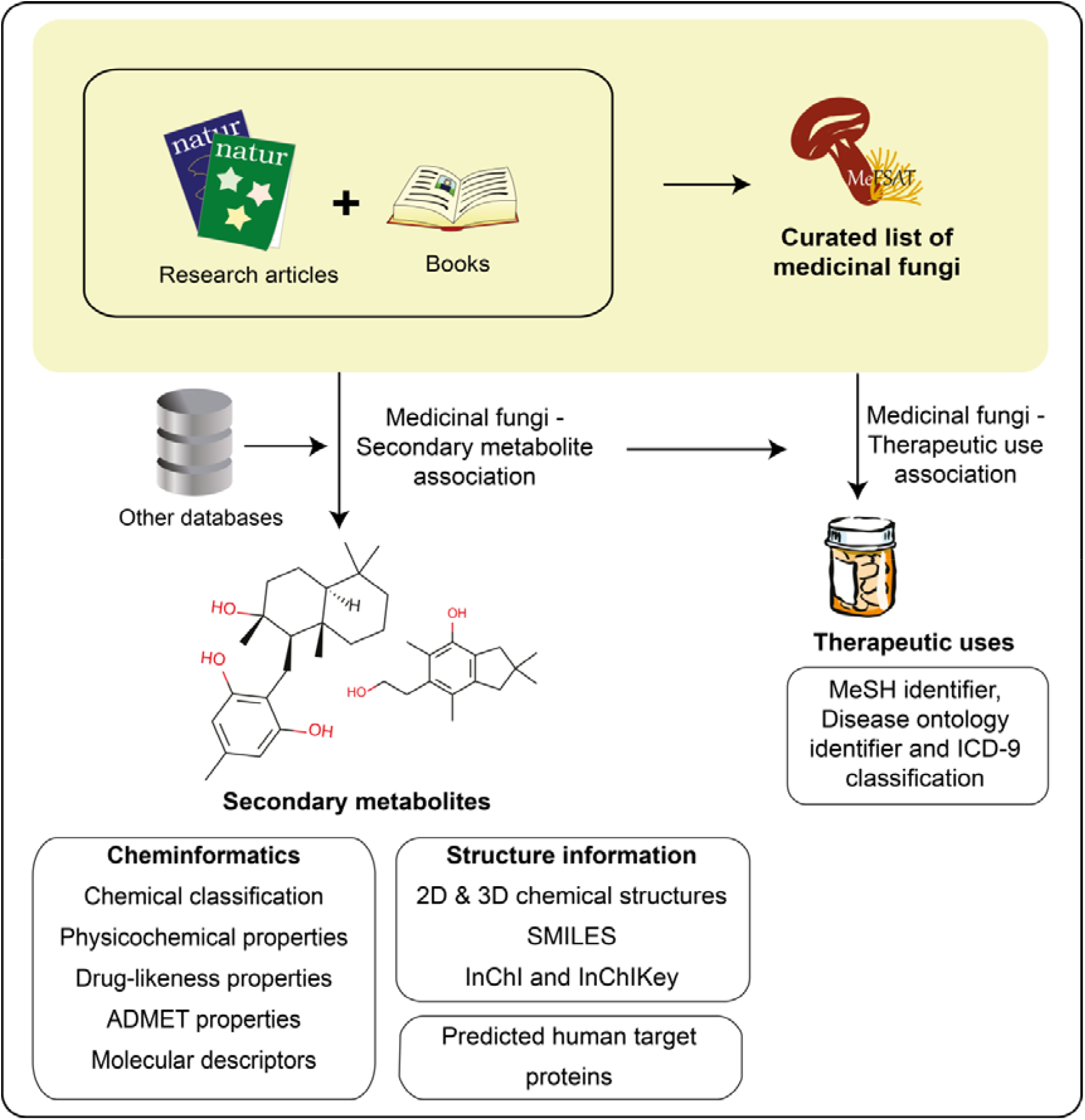
Schematic overview of the workflow to construct the MeFSAT database. Briefly, we compiled a curated list of medicinal fungi from the published literature. Next, we mined the published literature to compile secondary metabolites produced by different medicinal fungi. For the manually curated non-redundant list of secondary metabolites produced by medicinal fungi, we have compiled the chemical structures and employed cheminformatics tools to compute their physicochemical, drug-likeness and ADMET properties. Subsequently, we have compiled and curated information on the therapeutic uses of medicinal fungi from the published literature.

## Methods

### Workflow for the compilation and curation of secondary metabolites of medicinal fungi

Fungal secondary metabolites are a valuable chemical space of diverse natural products with several applications including in drug discovery^31^. **Me**dicinal **F**ungi **S**econdary metabolites **A**nd **T**herapeutics (MeFSAT) is a manually curated database that compiles information on secondary metabolites and reported therapeutic uses of medicinal fungi from published research articles and specialized books on the subject. In the following subsections, we provide an overview of the steps involved in the construction of MeFSAT database (Fig. 1).

### Curated list of medicinal fungi

The first step in the database construction workflow involved the compilation of a curated list of medicinal fungi from published literature. For this purpose, we performed an extensive PubMed^**32**^ search using the query “Medicinal fungi” OR “Medicinal mushroom”, and this keyword search last performed on 3 May 2020 led to the retrieval of 1220 published research articles. Apart from research articles, we also curated information on medicinal fungi from books^**17–21**^ on the topic. In the first-pass, we obtained a list of 354 fungi names with medicinal use from published literature consisting of research articles and books. Since the use of synonymous fungal names is common in the published literature, we next mapped the 354 fungal names with medicinal use to their accepted names using two resources namely, Catalogue of Life: 2019 Annual Checklist^**33**^ and Mycobank^**34**^. In the end, this mapping to accepted names led to a non-redundant list of 253 medicinal fungi.

### Secondary metabolites of medicinal fungi

The second step in our database construction workflow involved the search for secondary metabolites of 253 medicinal fungi in the published literature. From research articles and books^17–21,35^, we were able to gather information on 1139 secondary metabolites with evidence of being produced by at least one of 145 medicinal fungi. Further, we were able to gather additional information on 892 and 7 secondary metabolites with evidence of being produced by at least one of 121 and 5 medicinal fungi from two microbial natural product databases, namely, NPATLAS^15^ and novel Antibiotics database (http://www.antibiotics.or.jp/journal/database/database-top.htm), respectively. Overall, we were able to gather chemical names of 2038 secondary metabolites with evidence of being produced by at least one of 188 medicinal fungi from above-mentioned sources.

### Non-redundant *in silico* library of secondary metabolites of medicinal fungi

The primary objective of MeFSAT is to build a non-redundant curated resource of secondary metabolites of medicinal fungi along with information on their two-dimensional (2D) and three-dimensional (3D) chemical structure. Towards this objective, the compiled information on chemical names of secondary metabolites of medicinal fungi was evaluated to create a structurally non-redundant *in silico* chemical library. Specifically, the chemical names of the compiled secondary metabolites were mapped to chemical identifiers employed by standard chemical databases, namely, PubChem^36^, ChemIDplus (https://chem.nlm.nih.gov/chemidplus/), Chemspider^37^ and NPATLAS^15^. In case, we were unable to map a secondary metabolite to an identifier in at least one of the above-mentioned chemical databases, the chemical structure of the secondary metabolite was manually drawn from the corresponding published research article. Following the above-mentioned steps, we were able to obtain the chemical structure information for 1991 secondary metabolites which have evidence of being produced by at least one of 184 medicinal fungi. Note that a few secondary metabolites were omitted from further consideration as we were unable to either map them to an identifier or obtain their chemical structure from published literature.

Thereafter, we used an in-house Python script which employs Tanimoto coefficient^38^ (Tc) to determine chemical similarity between secondary metabolites. To create a non-redundant chemical library, we have merged compiled secondary metabolites from published literature if they were determined to be identical based on their chemical structure. Importantly, our manual curation effort while creating a non-redundant database of secondary metabolites has also taken into due consideration the stereochemistry of the chemical structures. Finally, this effort has led to a non-redundant set of 1830 secondary metabolites in MeFSAT database with literature evidence of being produced by at least one of 184 medicinal fungi.

### Annotation of secondary metabolites of medicinal fungi

For the 1830 secondary metabolites in MeFSAT database, the 2D chemical structures were saved in SDF, MOL and MOL2 file formats using OpenBabel^25^ while their 3D chemical structures were manually retrieved from PubChem^36^ if available. For the remaining secondary metabolites whose 3D structures are not available in PubChem, the 3D structures were generated using RDKit^25^ (http://www.rdkit.org/) by embedding the molecule using ETKDG method^39^ followed by energy minimization using MMFF94 force field^40^. The 3D structures for secondary metabolites were saved in SDF, MOL, MOL2, PDB and PDBQT file formats using OpenBabel^27^. Apart from the 2D and 3D structure information, the SMILES, InChI and InChIKey of the secondary metabolites were also generated using OpenBabel^27^. Moreover, the secondary metabolites in MeFSAT were hierarchically classified into chemical Kingdom, chemical SuperClass, chemical Class and chemical SubClass using ClassyFire^28^ (http://classyfire.wishartlab.com/).

The basic physicochemical properties of the secondary metabolites in MeFSAT database were computed using RDKit^25^ and SwissADME^26^ (http://www.swissadme.ch/). To assess the drug-likeness of the secondary metabolites in MeFSAT, we have computed multiple scoring schemes and properties namely, Lipinski’s rule of five (RO5)^41^, Ghose filter^42^, Veber filter^43^, Egan filter^44^, Pfizer’s 3/75 filter^45^, GlaxoSmithKline’s (GSK) 4/400^46^, Number of Leadlikeness violations^47^ and weighted quantitative estimate of drug-likeness (QEDw)^48^ using RDKit^25^ and SwissADME^26^. The assessment of Absorption, Distribution, Metabolism, Excretion and Toxicity (ADMET) properties is essential in the drug discovery pipeline. We have used SwissADME^26^ to predict the ADMET properties of the secondary metabolites in MeFSAT. Finally, we have computed 1875 (2D and 3D) molecular descriptors for each secondary metabolite in MeFSAT using PaDEL^29^ (http://padel.nus.edu.sg/software/padeldescriptor). These molecular descriptors can be broadly categorized into different classes such as chemical composition, topology, 3D shape and functionality.

### Genome sequences of medicinal fungi

For the 184 medicinal fungi that have secondary metabolite information is MeFSAT, we have gathered their genome sequencing status from the Joint Genome Institute (JGI) portal^49^ (https://genome.jgi.doe.gov/) and the National Center for Biotechnology Information (NCBI)^32^ (https://www.ncbi.nlm.nih.gov/genome). In addition, we have compiled information from published literature on the traditional system of medicine in which these medicinal fungi are used.

### Therapeutic uses of medicinal fungi

The next step in the database construction workflow involved the compilation of therapeutic uses of medicinal fungi from published literature including specialized books^20,21^. Note that the compiled therapeutic uses are based on available information on the use of medicinal fungi to treat human diseases. Notably, we have manually curated the compiled therapeutic use terms from various literature sources to create a non-redundant list of standardized therapeutic use terms for medicinal fungi in MeFSAT by mapping the therapeutic use terms to standard identifiers from Medical Subject Headings (MeSH)^50^, Disease Ontology^51^ and ICD-9-CM chapters^52^. In sum, this effort has led to a non-redundant list of 149 standardized therapeutic use terms in MeFSAT that are associated with different medicinal fungi.

### Predicted human target proteins of secondary metabolites

Lastly, we have compiled the predicted human target proteins of secondary metabolites in MeFSAT from the STITCH^30^ database (http://stitch.embl.de/). To date, STITCH^30^ is the largest resource on predicted interactions between chemicals and their target proteins. In MeFSAT database, we have included only high confidence interactions between secondary metabolites and human target proteins that have a combined STITCH^30^ score ≥ 700. Further, we have also mapped the genes corresponding to predicted human target proteins of secondary metabolites from STITCH^30^ to their respective HUGO Gene Nomenclature Committee (HGNC) symbols^53^.

### Comparison of physicochemical properties of fungal secondary metabolites with other small molecule collections

We have compared the physicochemical properties of secondary metabolites in MeFSAT with three other small molecule collections studied by Clemons *et al*^54^. The three small molecule libraries are: (a) a library of commercial compounds (CC) containing 6152 representative small molecules from commercial sources, (b) a library of diversity-oriented synthesis compounds (DC’) containing 5963 small molecules synthesized by the academic community, and (c) a library of natural products (NP) containing 2477 small molecules from various natural sources including microbes and plants. Note that the set of 1830 secondary metabolites in MeFSAT is also a natural product library, however, there is only a tiny overlap of 20 small molecules between NP library of Clemons *et al*^54^ and secondary metabolites in our database. Further, the computation of physicochemical properties failed for 3 small molecules each in CC and DC’ libraries of Clemons *et al*^54^, and thus, we have omitted them from later analysis.

### Chemical similarity of secondary metabolites

Tanimoto coefficient^38^ (Tc) is a widely used metric to compute chemical structure similarity^55^, and we have used Tc based on Extended Circular Fingerprints (ECFP4)^56^ to compute the structure similarity between secondary metabolites in MeFSAT and small molecule drugs approved by US FDA. For this purpose, the structures of FDA approved drugs were retrieved from DrugBank^57^. Note that the computed Tc value between any two molecules has a range between 0 and 1, wherein 0 represents little or no structure similarity and 1 represents very high or exact structure similarity. Based on a previous study by Jaisal *et al*^58^, we have chosen the cutoff of Tc ≥ 0.5 to decide if a given pair of chemicals have significant structure similarity.

### Chemical similarity network

We have constructed chemical similarity networks (CSNs) wherein nodes are chemicals and edges between pairs of chemicals signify high chemical similarity. For this construction, Tc based on ECFP4^56^ gives the extent of similarity between two chemicals, and thus, the weights of edges between nodes in the network. In the CSN, we only retain edges between two chemicals if the Tc between them is ≥ 0.5. We have constructed two CSNs here that correspond to 1830 secondary metabolites in MeFSAT and the subset of 228 drug-like secondary metabolites in MeFSAT, respectively. Further, these CSNs were visualized using Cytoscape^59^.

### Maximum common substructure

Maximum Common Substructure (MCS) of two or more chemical structures is the largest common substructure that is present in them. MCS has many applications including in chemical similarity search and hierarchical clustering of chemicals^60^. We have used RDKit^25^ to identify the MCSs for sets of secondary metabolites that form different clusters in the CSN comprising of 228 drug-like secondary metabolites in MeFSAT. Note that the MCSs were only computed for the 10 largest connected components (or clusters) in the CSN of drug-like secondary metabolites. Further, the identified MCSs were visualized using the online web-server SMARTSview^61^ (https://smartsview.zbh.uni-hamburg.de/).

### Web interface and database management system

MeFSAT database compiles information mainly on two types of association, (a) medicinal fungi and their secondary metabolites, and (b) medicinal fungi and their therapeutic uses. In addition, MeFSAT compiles detailed information for secondary metabolites such as their 2D and 3D chemical structure, chemical classification, physicochemical properties, drug-likeness properties, predicted ADMET properties, molecular descriptors and predicted human target proteins. MeFSAT is openly accessible at: https://cb.imsc.res.in/mefsat/.

The compiled information in MeFSAT is stored in a SQL database created using the open-source relational database management system MariaDB (https://mariadb.org/). The web interface of MeFSAT was created using the free and open-source CSS framework Bootstrap 4.1.3 (https://getbootstrap.com/docs/4.0/getting-started/introduction/), with custom HTML, PHP (http://php.net/), CSS, JavaScript and jQuery (https://jquery.com/) scripts. To render visualizations in the web interface, we have used Cytoscape.js (http://js.cytoscape.org/) and Google Charts (https://developers.google.com/chart). The MeFSAT website is hosted on an Apache server (https://httpd.apache.org/) running on Debian 9.13 Linux operating system.

## Results and discussion

### MeFSAT database and its web interface

In this work, we have built a natural product database, namely, **Me**dicinal **F**ungi **S**econdary metabolites **A**nd **T**herapeutics (MeFSAT), which contains manually curated information on secondary metabolites and therapeutic uses of medicinal fungi from published literature (Methods; Fig. 1). The database has a modern and intuitive web interface that enables easy access 11 to the curated information (Fig. 2). MeFSAT is openly accessible at: https://cb.imsc.res.in/mefsat/.

**Figure 2.**
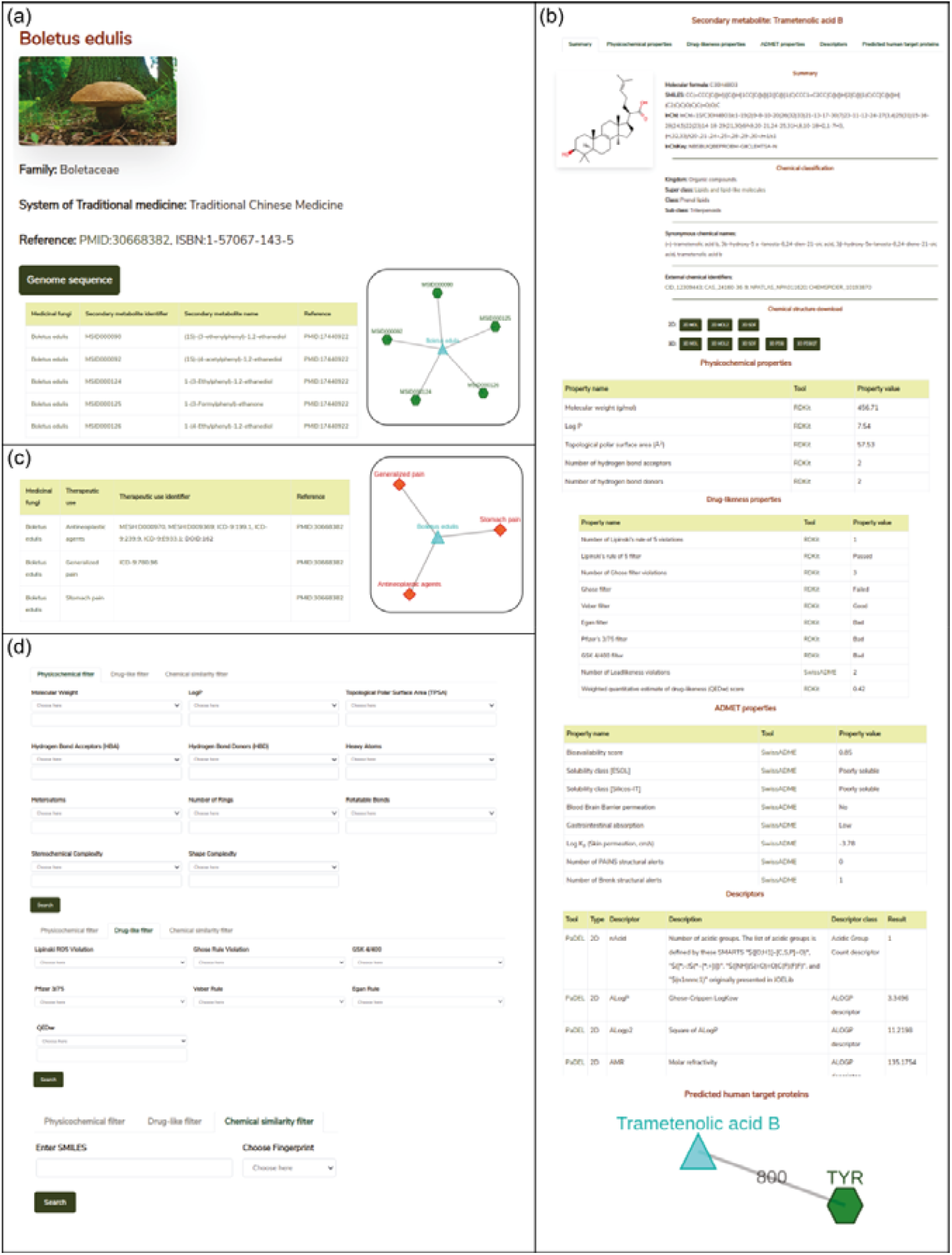
Web interface of the MeFSAT database. **(a)** Snapshot of the result of a standard query for secondary metabolites of a medicinal fungus. The example shows the secondary metabolites for the fungus *Boletus edulis*. **(b)** Snapshot of the detailed information page for a secondary metabolite which gives its 2D and 3D chemical structure, physicochemical properties, drug-likeness properties, predicted ADMET properties, molecular descriptors and predicted human target proteins. From this page, users can download the 2D and 3D structure of the secondary metabolite in SDF, MOL, MOL2, PDB or PDBQT file formats. The example shows information for the secondary metabolite Trametenolic acid B. **(c)** Snapshot of the result of a standard query for therapeutic uses of a medicinal fungus. The example shows the therapeutic uses for the fungus *Boletus edulis*. **(d)** Snapshot of the advanced search options which enable users to filter secondary metabolites based on their physicochemical properties or drug-likeness properties or chemical similarity with a query chemical structure in SMILES format.

The MeFSAT web interface enables users to retrieve manually curated associations between medicinal fungi and secondary metabolites or therapeutic uses by querying for either (a) scientific names of medicinal fungi, (b) secondary metabolite identifier, (c) secondary metabolite name, or (d) therapeutic use terms (Fig. 2). The query result is displayed as a table of associations with relevant literature references. In resultant table obtained after the search for secondary metabolite associations of medicinal fungi, the users can click on a specific medicinal fungi name that will redirect them to a dedicated page containing all secondary metabolite associations for the specific fungi (Fig. 2a). The users can also view the detailed information page for each secondary metabolite by clicking the secondary metabolite identifiers in the above-mentioned table. The detailed information page for a secondary metabolite provides a summary of the chemical structure, external database identifiers, synonymous names, chemical classification, 2D and 3D chemical structure in different file formats, physicochemical properties, drug-likeness properties, predicted ADMET properties, molecular descriptors and predicted human target proteins (Methods; Fig. 2b).

In resultant table obtained after the search for therapeutic use associations of medicinal fungi, the users can click on a specific medicinal fungi name that will redirect them to a dedicated page containing therapeutic uses of the specific fungi which were curated from published literature along with the identifiers for therapeutic use terms from MeSH^50^, Disease ontology^51^ and ICD-9-CM chapters^52^ (Fig. 2c). Further, by clicking the therapeutic use terms in the above-mentioned table, the users can view the list of medicinal fungi that have a specific therapeutic use.

Advanced search options in MeFSAT web interface enable users to filter the secondary metabolites by either: (a) physicochemical properties, (b) drug-likeness properties, or (c) chemical structure similarity (Fig 2d). Specifically, the physicochemical filter tab enables users to retrieve secondary metabolites with desired physicochemical properties. The drug-likeness filter tab enables users to select secondary metabolites that pass or fail multiple drug-likeness scoring schemes. Lastly, the chemical similarity filter enables users to search for 10 secondary metabolites within MeFSAT that have the highest structural similarity to a query chemical compound entered in SMILES format. The results from these advanced search options are rendered as tables wherein secondary metabolites can be sorted based on the chosen properties.

### Curated information on medicinal fungi, their secondary metabolites and therapeutic uses

The MeFSAT database compiles manually curated information on 1830 secondary metabolites produced by at least one of 184 medicinal fungi (Methods; ESI Table S1). For this curated list of 184 medicinal fungi in MeFSAT, we have compiled information on their taxonomic family, genome sequencing status, and usage in different systems of traditional medicine (Methods). Interestingly, 54 out of the 184 medicinal fungi in MeFSAT are used in traditional Chinese medicine to treat human ailments. Further, the 184 medicinal fungi are distributed across 48 taxonomic families of which the 5 families Polyporaceae, Ganodermataceae, Agaricaceae, Hymenochaetaceae and Pleurotaceae have 24, 20, 18, 13 and 10 medicinal fungi, respectively, in MeFSAT database (ESI Table S1).

There are 2127 medicinal fungi - secondary metabolite associations in MeFSAT database which encompass 184 medicinal fungi and 1830 secondary metabolites (Methods; Fig. 1). These 1830 secondary metabolites are distributed across 13 chemical SuperClasses as computed using ClassyFire^28^ (Methods, Fig. 3a). Notably, more than 60% of the secondary metabolites in 13 MeFSAT belong to the chemical SuperClass ‘Lipids and lipid-like molecules’. Other chemical SuperClasses enriched in secondary metabolites from MeFSAT include ‘Organoheterocyclic compounds’ (14%), ‘Organic oxygen compounds’ (7%) and ‘Benzenoids’ (7%) (Fig. 3a). Among the 184 medicinal fungi, *Ganoderma lucidum* has the highest number (277) of secondary metabolite associations, followed by *Ganoderma applanatum* with 131 secondary metabolite associations, and *Hericium erinaceus* with 104 secondary metabolite associations.

**Figure 3.**
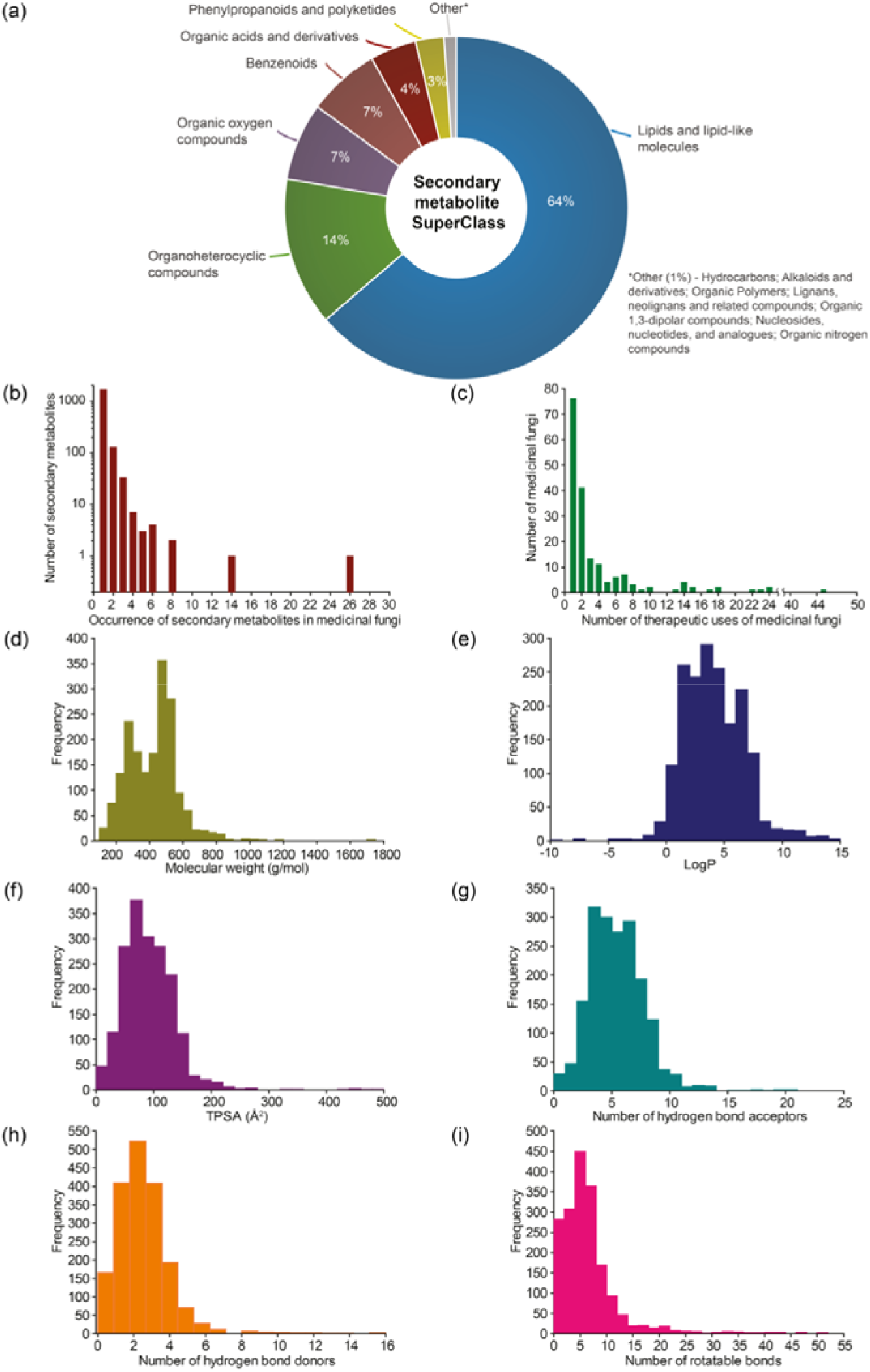
Basic statistics for medicinal fungi, their secondary metabolites and therapeutic uses in MeFSAT database. **(a)** Pie chart shows the distribution of the secondary metabolites in MeFSAT across chemical SuperClasses obtained from ClassyFire^21^ . **(b)** Histogram of the number of medicinal fungi with literature evidence of producing a given secondary metabolite in MeFSAT database. **(c)** Histogram of the number of therapeutic uses per medicinal fungi in MeFSAT database. **(d–i)** Histogram of the distribution of molecular weight (g/mol), LogP, topological polar surface area (TPSA) (Å^2^), number of hydrogen bond acceptors, number of hydrogen bond donors, and number of rotatable bonds, for the secondary metabolites in MeFSAT database.

In Fig. 3b, we show the histogram of the occurrence of secondary metabolites across 184 medicinal fungi in MeFSAT database. It is seen that the majority of the secondary metabolites (1779) in MeFSAT database have published literature evidence of being produced by less than 3 medicinal fungi. Further, only two secondary metabolites in MeFSAT database have published literature evidence of being produced by more than 10 medicinal fungi; these metabolites are L-ergothioneine (MSID001103) with evidence from 26 medicinal fungi and Ergosterol peroxide (MSID000285) with evidence from 14 medical fungi. MeFSAT database also compiles 2036 predicted interactions between secondary metabolites and their human target proteins from the STITCH^30^ database, and these interactions encompass 54 secondary metabolites and 1003 human proteins (Methods).

There are 689 medicinal fungi - therapeutic use associations in MeFSAT database which encompass 179 medicinal fungi and 149 therapeutic uses (Methods; Fig. 1). Fig. 3c shows the histogram of the number of therapeutic uses per medicinal fungi as compiled in MeFSAT database. It is seen that the majority of the medicinal fungi (162) in MeFSAT have less than 10 reported therapeutic uses, while 5 medicinal fungi have more than 20 reported therapeutic uses in published literature. *Ganoderma lucidum* has the highest number (45) of reported therapeutic uses, followed by *Hericium erinaceus* (24), *Lignosus rhinocerus* (24), *Antrodia cinnamomea* (23) and *Tropicoporus linteus* (22).

### Physicochemical properties of secondary metabolites in MeFSAT database and comparison with other small molecule collections

For the 1830 secondary metabolites in MeFSAT database, we have computed several physicochemical properties (Methods). In Fig. 3d-i, we show the distribution of six physicochemical properties namely, molecular weight, LogP, topological polar surface area (TPSA), number of hydrogen bond acceptors, number of hydrogen bond donors, and number of rotatable bonds across the 1830 secondary metabolites in MeFSAT database.

Previously, Clemons *et al*^54^ have shown that two size-independent metrics namely, stereochemical complexity and shape complexity, are excellent predictors of target protein binding specificity of small molecules (Methods). Stereochemical complexity measures the ratio of the number of chiral carbon atoms to the total number of carbon atoms in a molecule, whereas shape complexity is the ratio of the number of sp^3^ hybridized carbon atoms to the total number of sp^2^ and sp^3^ hybridized carbon atoms in a molecule^54^. Specifically, Clemons *et al*^54^ have correlated the stereochemical and shape complexity of small molecules with their target protein binding specificity across three representative small molecule collections namely, CC, DC’ and NP (Methods). Small molecules in NP collection were found to have higher stereochemical and shape complexity in comparison with those in DC’ or CC collection, and moreover, small molecules in NP collection were found to be more specific binders of target proteins with low fraction of promiscuous binders in comparison with those in DC’ or CC collection^54^. In other words, natural products^14,54^ were found to have higher stereochemical and shape complexity while being specific binders of target proteins.

Here, we have computed and compared the stereochemical complexity and shape complexity of the 1830 secondary metabolites in MeFSAT with those of small molecules in CC, DC’ and NP collections (Methods; Fig 4a-b). Interestingly, we find that the mean and median of the stereochemical complexity or shape complexity of secondary metabolites in MeFSAT are closer to the NP collection than DC’ or CC collections. This suggests that secondary metabolites in MeFSAT are more likely to be specific binders of target proteins than promiscuous binders. Moreover, apart from stereochemical and shape complexity, we have also compared the mean and median of six other physicochemical properties namely, molecular weight, LogP, TPSA, number of hydrogen bond acceptors, number of hydrogen bond donors and number of rotatable bonds, for the 1830 secondary metabolites in MeFSAT with those for small molecules in NP, DC’ and CC collections (Fig. 4c).

**Figure 4.**
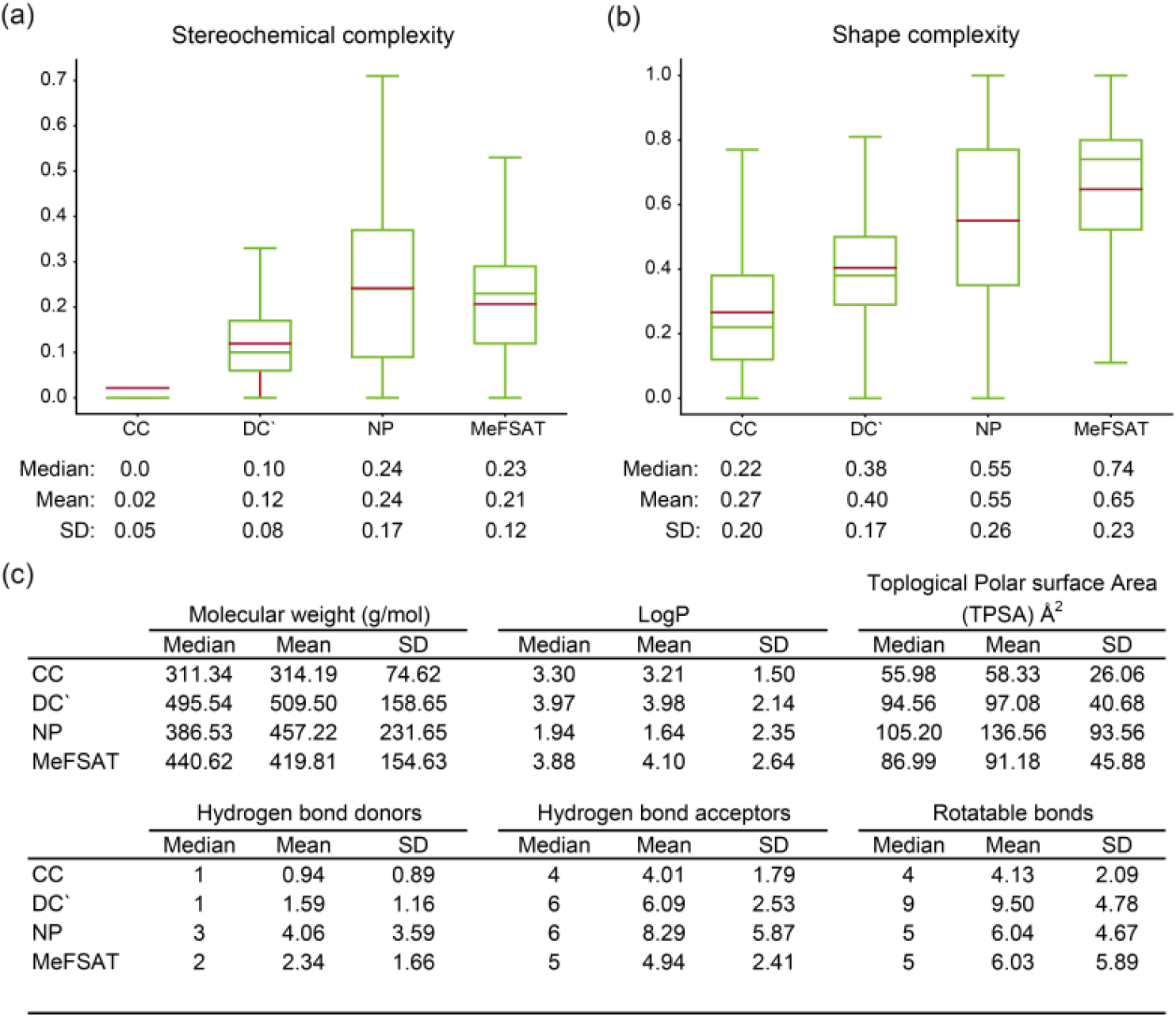
Comparison of the physicochemical properties of secondary metabolites in MeFSAT with other small molecule collections. (a) Box plot shows the distribution of the stereochemical complexity of the small molecule collections CC, DC’, NP and MeFSAT secondary metabolites. The median, mean and standard deviation (SD) of the stereochemical complexity for each small molecule collection is shown below the box plot. (b) Box plot shows the distribution of the shape complexity of the small molecule collections CC, DC’, NP and MeFSAT secondary metabolites. The median, mean and SD of the shape complexity for each small molecule collection is shown below the box plot. Note the lower end of the box shows the first quartile, upper end of the box shows the third quartile, green line shows the median and brown line shows the mean of the distribution of stereochemical complexity or shape complexity in the two box plots. (c) Median, mean and SD of six physicochemical properties, namely, molecular weight, logP, topological polar surface area (TPSA), number of hydrogen bond donors, number of hydrogen bond acceptors, and number of rotatable bonds for the small molecule collections CC, DC’, NP, and MeFSAT secondary metabolites.

### Drug-like secondary metabolites of medicinal fungi

Natural products have directly or indirectly contributed to the discovery of ∼35% of the small molecules drugs approved by the US FDA till 2014^6^. To date, several scoring schemes or rules have been proposed to assess the drug-likeness of small molecules^62^. Here, we have used six scoring schemes namely, Lipinski’s RO5^41^, Ghose filter^42^, Veber filter^43^, Egan filter^44^, Pfizer’s 3/75 filter^45^ and GSK 4/400^46^ to access the drug-likeness of 1830 secondary metabolites in MeFSAT (Methods). Notably, we have identified a subset of 228 secondary metabolites (∼12%) in MeFSAT to be drug-like, and these metabolites have passed all the six scoring schemes mentioned above. In Fig. 5a, we show the number of secondary metabolites in MeFSAT that pass different combinations of above-mentioned six scoring schemes. It is important to highlight that several natural products that failed to pass drug-likeness scores have been successfully developed into drugs^63^. Therefore, we expect a higher fraction of secondary metabolites in MeFSAT, greater 16 than the ∼12% metabolites that pass the six scoring schemes, have the potential to be developed into drugs. Fig. 5b shows the chemical classification of the 228 drug-like secondary metabolites that pass the six scoring schemes (Methods). The 228 drug-like secondary metabolites in MeFSAT were distributed across 8 chemical SuperClasses, with more than 30% classified as ‘Organoheterocyclic compounds’. Furthermore, based on the computation of the QEDw metric, we find that 19 out of 228 drug-like secondary metabolites in MeFSAT have a high QEDw value of > 0.80 (Fig. 5c). This further highlights the potential of the MeFSAT chemical space for drug discovery.

**Figure 5.**
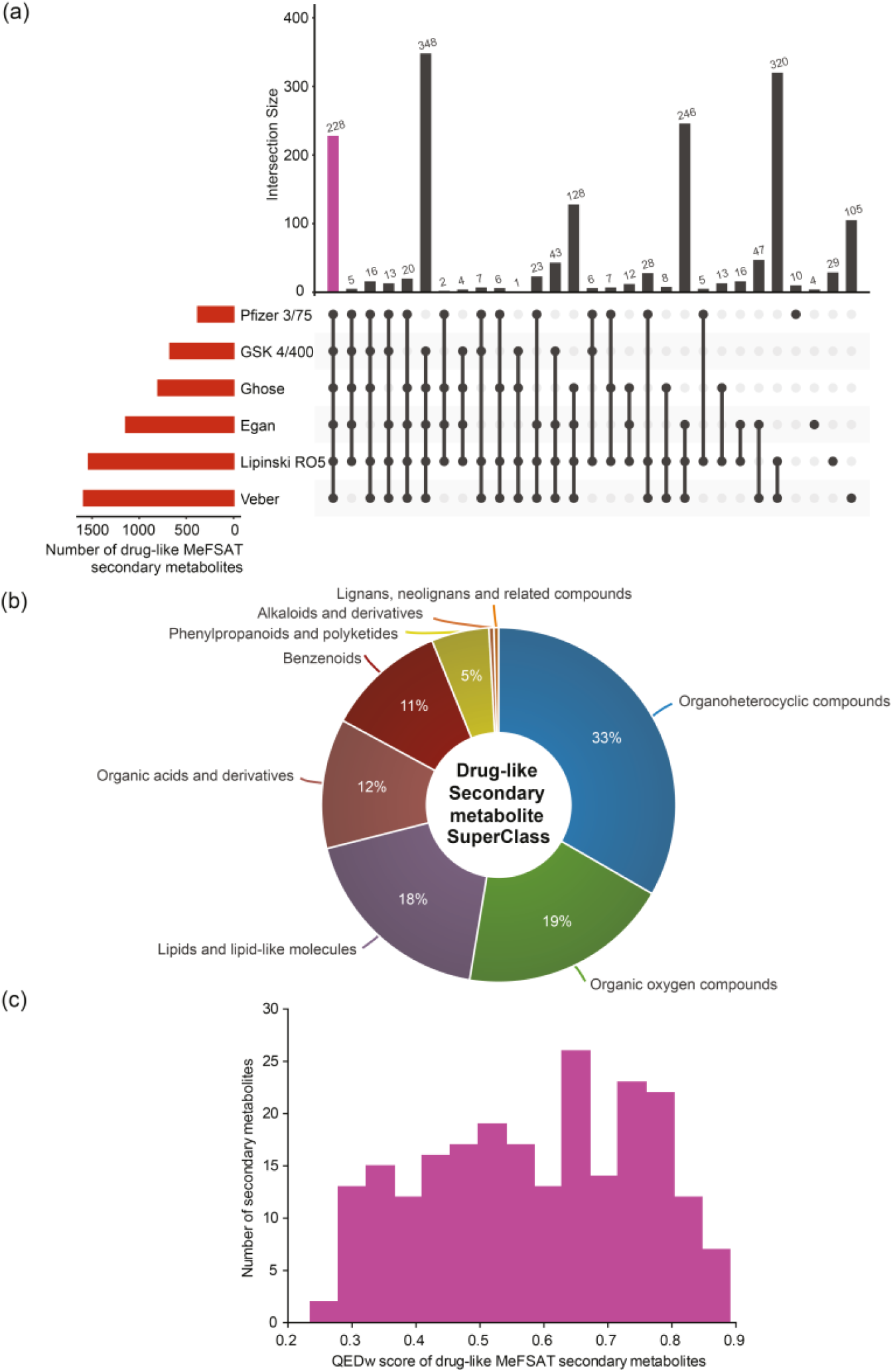
Drug-likeness analysis of the secondary metabolites in MeFSAT database. **(a)** Evaluation of drug-likeliness of secondary metabolites based on multiple scores. The horizontal bar plot shows the number of secondary metabolites in MeFSAT database that satisfy different drug-likeness scoring schemes (Methods). The vertical bar plot shows the overlap between sets of secondary metabolites that satisfy different drug-likeness scoring schemes. The pink bar in the vertical plot gives the 228 secondary metabolites that satisfy all the 6 drug-likeness scoring schemes. This plot was generated using the UpSetR package^63^. **(b)** Classification of the 228 drug-like secondary metabolites into chemical SuperClasses obtained from ClassyFire^21^. **(c)** Distribution of the QEDw scores for the 228 drug-like secondary metabolites that satisfy all the 6 drug-likeness scoring schemes.

### Chemical similarity networks of secondary metabolites

Chemical similarity networks (CSNs) can facilitate visualization and exploration of the structural diversity in a chemical library, and thus, enable better selection of lead compounds from a chemical space for drug development. Here, we have constructed two CSNs (Methods) wherein the first corresponds to 1830 secondary metabolites (ESI Fig. S1) while the second to 228 drug-like secondary metabolites (Fig. 6a) in MeFSAT database. In the CSNs, the nodes representing secondary metabolites are colored in green if they are similar to any of the FDA approved drugs, else they are colored in pink (Methods; Fig. 6a; ESI Fig. S1). Moreover, the thicknesses of edges in CSNs reflect the chemical similarity between pairs of secondary metabolites connected by them (Methods).

**Figure 6.**
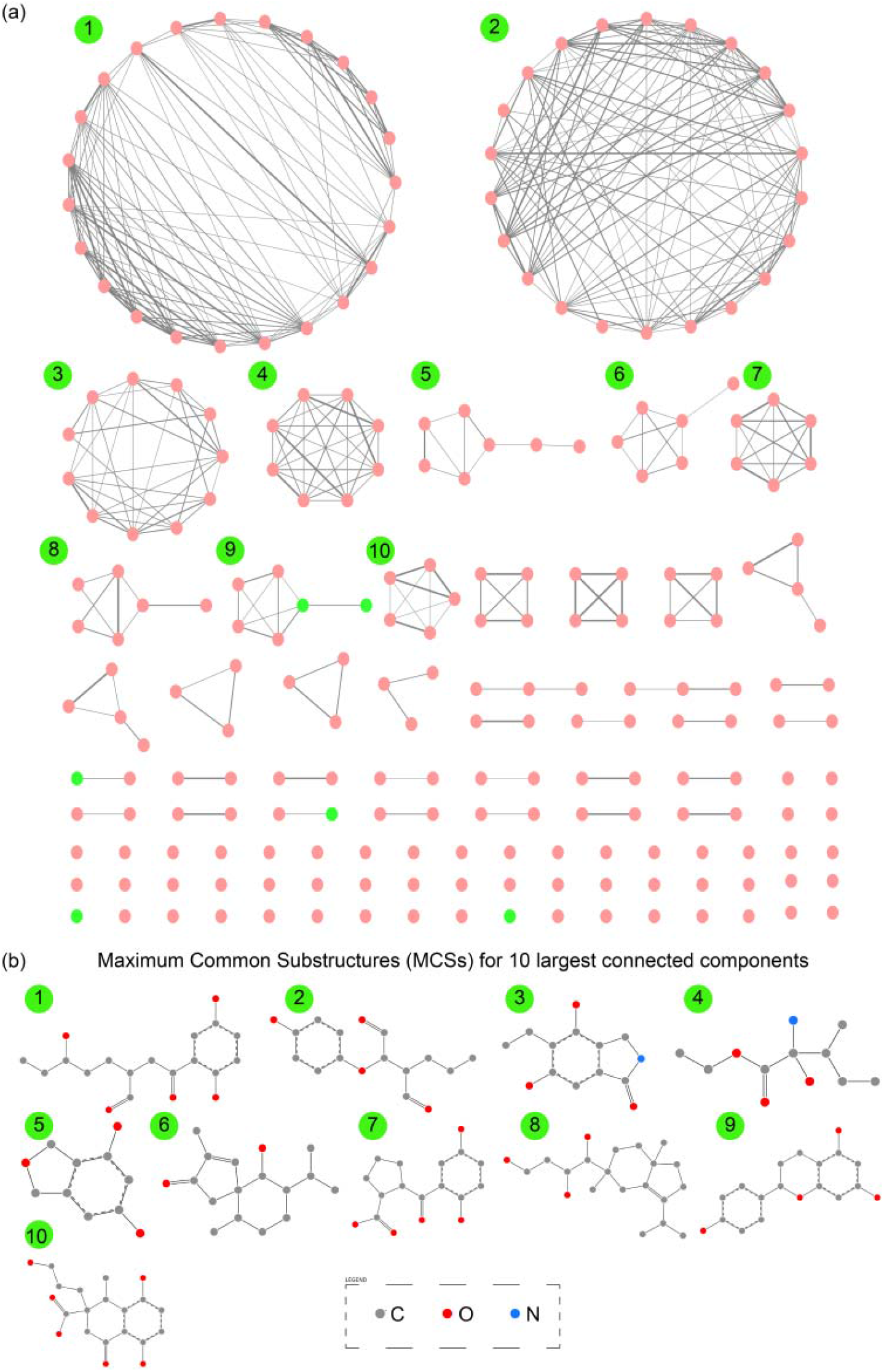
**(a)** Chemical similarity network (CSN) of 228 drug-like secondary metabolites in MeFSAT database. Here, the node color is green if the corresponding secondary metabolite is similar to any of the FDA approved drugs else the node color is pink. Edge thickness is proportional to the computed structural similarity between pairs of secondary metabolites. **(b)** Maximum Common Substructures (MCSs) for 10 largest connected components in the CSN of drug-like secondary metabolites. For this figure, the visualization of the SMARTS was generated using SMARTSview^59^.

Specifically, we find that 82 of 1830 secondary metabolites and 6 of 228 drug-like secondary metabolites in MeFSAT database have high similarity to at least one of the FDA approved drugs (Methods). The two CSNs contain multiple disconnected clusters and several isolated nodes which underscore the rich structural diversity in the chemical space of secondary metabolites in MeFSAT database (Fig. 6a; ESI Fig. S1). Specifically, the CSN of 1830 secondary metabolites 17 has 335 connected components of which 206 are isolated nodes (ESI Fig. S1). Similarly, the CSN of 228 drug-like secondary metabolites has 94 connected components of which 55 are isolated nodes (Fig. 6a). Moreover, graph density which is a measure of the fraction of all possible edges that are realized in the network, is found to be 0.01 and 0.02, respectively, for the CSNs of 1830 secondary metabolites and 228 drug-like secondary metabolites, respectively. This overly sparse nature of the two CSNs further underscores the structural diversity of the chemical space of secondary metabolites in MeFSAT database.

Finally, we have also computed the maximum common substructures (MCSs) for the 10 largest connected components within the CSN of 228 drug-like secondary metabolites (Methods; Fig. 6b; ESI Table S2). The computed MCSs display the unique structural motifs underlying the 10 largest connected components in the CSN of drug-like secondary metabolites (Fig. 6b).

## Conclusions

There is immense interest in tapping the diverse chemical space of fungal secondary metabolites for drug discovery. For centuries, medicinal fungi including mushrooms have been used in many systems of traditional medicine to treat human ailments. Since the therapeutic action of medicinal fungi is likely due to their novel secondary metabolites, a curated database compiling information on secondary metabolites and therapeutic uses of medicinal fungi will be a valuable resource for computer-aided drug discovery. Therefore, we have built the first dedicated resource MeFSAT compiling information on secondary metabolites and therapeutic uses of medicinal fungi from published literature. MeFSAT compiles manually curated information on 184 medicinal fungi, 1830 secondary metabolites and 149 therapeutic uses from published literature. For the non-redundant library of 1830 secondary metabolites, we have compiled information on their chemical structure, physicochemical properties, drug-likeness properties, predicted ADMET properties, molecular descriptors, and predicted human target proteins. MeFSAT database also compiles taxonomic information, genome sequencing status, system of traditional medicine and therapeutic uses of medicinal fungi.

After building the MeFSAT database, we have compared the distributions of physicochemical properties for the 1830 secondary metabolites with those for three other small molecule collections. Based on this comparative analysis, we find that the stereochemical complexity and shape complexity of secondary metabolites in MeFSAT are similar to those for other natural product libraries. Based on the previous work by Clemons *et al*^54^, one can also extrapolate that the secondary metabolites of medicinal fungi are likely to be specific binders of target proteins. Using multiple scoring schemes, we have also filtered a subset of 228 drug-like secondary metabolites within the MeFSAT database. Lastly, the construction and analysis of chemical similarity networks of secondary metabolites underscores the high structural diversity of the associated chemical space. In conclusion, MeFSAT is a curated resource of fungal natural products which will facilitate computational drug discovery.

## Supporting information

ESI Tables S1-S2

ESI Figure S1

## Author Contributions

**R.P. Vivek-Ananth:** Data curation, Formal analysis, Software, Visualization, Writing – original draft. **Ajaya Kumar Sahoo:** Data curation, Formal analysis, Software, Visualization, Writing – original draft. **Kavyaa Kumaravel:** Data curation, Writing – original draft. **Karthikeyan Mohanraj:** Software, Visualization. **Areejit Samal:** Conceptualization, Formal analysis, Supervision, Writing – review & editing.

## Acknowledgements

We thank B. Raveendra Reddy for computational support. AS acknowledges support from a Ramanujan fellowship (SB/S2/RJN-006/2014) from the Science and Engineering Research Board (SERB) India, the Department of Atomic Energy (DAE) India, and a Max Planck Partner Group in Mathematical Biology from the Max Planck Society Germany. The funders have no role in study design, data collection, data analysis, manuscript preparation or decision to publish.

## Electronic Supplementary Information (ESI) available

**Table S1**. The table gives the curated list of 184 medicinal fungi for which secondary metabolite information has been compiled from published literature in MeFSAT database. Cited references are provided in PMID, DOI and ISBN formats.

**Table S2**. Maximum common substructures (MCSs) for the 10 largest connected components in the CSN of drug-like secondary metabolites. The MCSs are listed in SMARTS format and were computed using RDKit.

**Figure S1**. Chemical similarity network (CSN) of 1830 secondary metabolites in MeFSAT database. Here, the node color is green if the corresponding secondary metabolite is similar to any of the FDA approved drugs else the node color is pink. Edge thickness is proportional to the computed structural similarity between pairs of secondary metabolites.

