## Supplementary material for "MeFSAT: A curated natural product database specific to secondary metabolites of medicinal fungi": ESI Figure S1

### Electronic Supplementary Information (ESI)

### Figure S1

### for

### MeFSAT: A curated natural product database specific to secondary metabolites of medicinal fungi

R. P. Vivek-Ananth[a,b,1], Ajaya Kumar Sahoo[a,b,1], Kavyaa Kumaravel[a,1], Karthikeyan Mohanraj[a,c], Areejit Samal[a,b,]*

[a] *The Institute of Mathematical Sciences (IMSc), Chennai 600113, India*

[b] *Homi Bhabha National Institute (HBNI), Mumbai 400094, India*

[c] *Institute for Clinical Chemistry and Laboratory Medicine, Technische Universität Dresden, Dresden 01307, Germany*

[1] R.P.V., A.K.S. and K.K. contributed equally to this work and should be considered as Joint-First authors

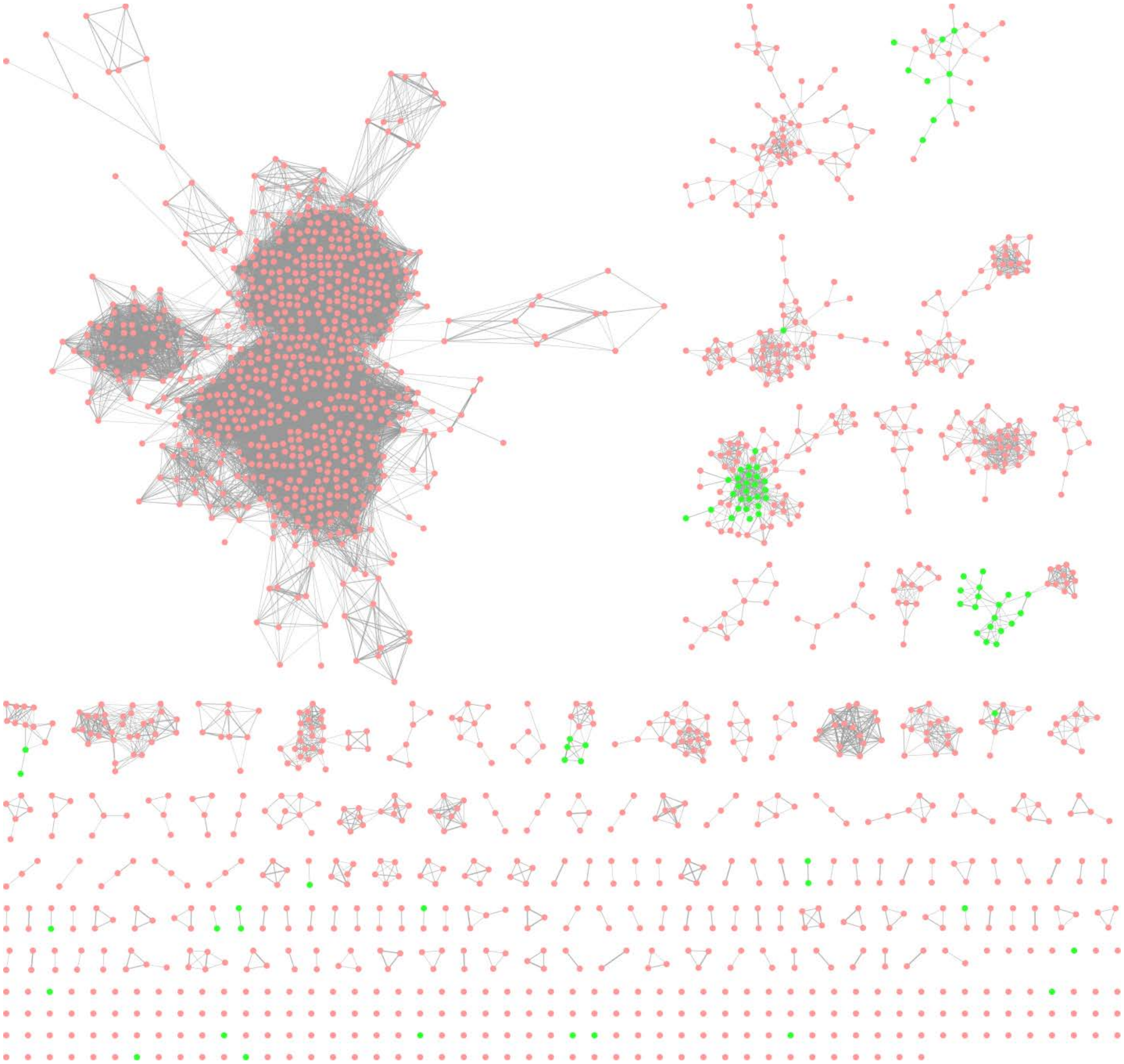

**Figure S1.** Chemical similarity network (CSN) of 1830 secondary metabolites in MeFSAT database. Here, the node color is green if the corresponding secondary metabolite is similar to any of the FDA approved drugs else the node color is pink. Edge thickness is proportional to the computed structural similarity between pairs of secondary metabolites.
